# An AI-ready compositional framework for mechanistic aging research and *in silico* intervention testing

**DOI:** 10.64898/2026.09.22.753641

**Authors:** Minja Belic, David Furman

**Affiliations:** Buck Institute for Research on Aging, Novato, CA, USA

## Abstract

Aging is a network-level phenomenon, with its hallmarks interacting through dense feedback loops across vastly different timescales. Decades of reductionist research have produced thousands of mechanistic models of narrow subsystems, but no straightforward way to integrate them into one comprehensive system. Consequently, whole-cell and multi-hallmark aging models remain rare, manually constructed, and relatively small, despite broad agreement on their importance. The emergence of autonomous research agents offers a way to distribute this modeling effort across humans and artificial intelligence and thereby greatly accelerate it, but only if the underlying substrate allows agents to readily compose and analyze models with built-in validation checks and modeling guidance, without extensive custom code.

We present *hallsim*, a JAX-native compositional simulation framework designed as the mechanistic substrate for agent-orchestrated research. It provides composability, end-to-end differentiability, GPU execution, and automated analysis and model-selection tools that facilitate composite model construction and fine tuning. The framework makes it easier to propose a mechanistic hypothesis, integrate it into an existing composite, reparametrize it against observed data, and batch-test it across different initial conditions. We demonstrate the co-simulation and in-silico perturbation of three independently published kinetic models spanning genomic instability, nutrient sensing, and proteostasis. The models are connected by four edges and calibrated against a public dataset. Additionally, we train a Neural ODE surrogate and compose it alongside the mechanistic modules in a hybrid composite, demonstrating that mechanistic and neural models can function as complementary components of the same system.

## 1. Introduction

Aging biology has revealed itself to be a complex-systems problem [1]. The widely accepted hallmarks of aging [2] with twelve interacting axes explicitly frame deterioration as a product of crosstalk rather than a failure of any one pathway. To what extent the biology of aging is emergent from its mechanistic constituents is unclear. Devising elaborate mechanistic models, simulating them at scale, and verifying against experimental observations might provide some insight into this question. In the past, the scientific community has predominantly adopted the reductionist view of biology which produced thousands of relevant mechanistic models for kinetic processes that explain the behavior of a narrow subsystem, but these models have largely been built by different research groups, correspond to different scopes, and are distributed in different formats (SBML, CellML, XPP, custom MATLAB, Python, Julia, and more). There have been rare attempts to combine some of the models relevant to aging into a larger composite system [3], [4], however this manual approach is time consuming and does not scale, which is also why there have only been a handful of whole-cell models despite the field’s agreement on this being an important goal [5].

The recent push to automate science and the emergence of multiple semi-autonomous artificial intelligence (AI) researchers (Kosmos [6], Biomni [7], and others) offers a new hope that the modelling effort can be distributed between human researchers and AI agents, and thus vastly accelerated, giving us perhaps an understanding of aging biology at the scale that wasn’t possible before. However, the available infrastructure is not sufficient to allow AI agents to keep combining the existing knowledge, training complementary neural based models from data where knowledge is lacking, and creating new insights in this domain. While established tools like Tellurium [8], and COPASI [9] are mature and heavily optimized, they lack the automatic differentiation needed to fit custom neural-based components and were not built for programmatic composition at scale. Vivarium [10] offers elegant architectural solutions for composition, but its execution model is CPU-bound and autodifferentiation is out of reach. JAX emerged as an ecosystem that solves this differentiability gap as demonstrated by SBMLtoODEjax [11], a Python library that converts SBML models to tunable JAX modules, and jaxkineticmodel [12] for gradient-based fitting of kinetic models, though it has yet to enter mainstream use. Model repositories like BioModels [13] standardize storage, BioSimulators

[14] and JWS Online [15] also provide a platform for individual simulation, but neither offers plug-and-play composition, one would have to write hundreds of lines of custom code to edit and rewire these models in order to run them together and reparametrize them to data. The result is that integrative, multi-hallmark mechanistic studies remain manual, brittle, and smallscale.

A parallel argument in the field claims this is the wrong problem to solve at all: foundation models trained on omics data will replace mechanistic modelling, and the manual integration problem will become irrelevant. We are skeptical for two reasons. First, foundation models trained on observational data interpolate within the support of their training distribution; they offer no first-principles basis for causal counterfactuals like geroprotective interventions. Emerging large perturbation models trained on single cell perturbSeq data partly address this limitation, although their performance relative to simple baselines remains mixed. Some evaluations find no consistent advantage for deep-learning methods [16], whereas others report improvements dependent on the prediction task [17]. These advances still do not eliminate the need for explicit dynamical models, since predicting a transcriptional response at a sampled endpoint does not establish the intervening trajectory or the response to a different intervention schedule. We anticipate that learning these dynamics will become an increasingly important focus of the field. Second, decades of mechanistic knowledge are encoded in the existing model corpus; abandoning it amounts to discarding a rare resource that has already been peer-reviewed. Enrichment of deep learning models with curated expert knowledge such as gene-gene interaction graphs was shown to improve perturbation prediction performance [17], [18]. This motivates a hybrid approach: mechanistic backbones for what we know causally, neural components (either transformers, Neural ODEs or other) for unknown modules. Alternatively, mechanistic models could potentially be used to constrain loss functions in the training of deep learning models in PINN style [19], [20].

This paper presents hallsim, a JAX-native compositional simulation framework that operationalizes this hybrid view. Composites from many published kinetic models wired into a single multi-process system form the mechanistic backbone, while neural modules for the biology we cannot yet write down are embedded as components of that same composite and can be trained jointly with it. We implement multiple features that facilitate model choice and composition, from searching literature, model and data repositories, through proposition of wiring targets, semantic validation of composition, bifurcation analysis, solver selection and identifiability analysis. Role-specific LLM prompts provide additional critiques of biological assumptions and mathematical formulation. We argue that the future will involve automated research through LLM orchestrated research agents, and hallsim is a substrate that such agents need to propose mechanistic hypotheses and eventually iteratively test them at scale with the help of lab automation.

## 2. Architecture

### 2.1 Composition

Hallsim’s composition solution is adapted from Vivarium [10] and reimplemented in JAX with performance-related adaptations and machine-friendly add-ons. Its purpose is to let a model be written once in isolation, and then combined with others by wiring, so that a mechanism published by one group can be joined to one published by another without touching either one’s code. This is operationalized through five object types: Process, Port, Topology, Store, and Composite. A Process is a self-contained model of one mechanism, such as a signaling module, a metabolic branch, or a coupling edge, implemented as an Equinox module [21]. A port is a connection point on a Process. It can be continuous, discrete, or event-driven and its role declares how the process interacts with the state behind it. An “evolved” port contributes additively, so that several processes writing to the same variable have their derivatives summed, an “exclusive” port claims sole ownership of a variable, an “input” port reads without contributing, and a “latched” port is written by discrete or event processes. Each port carries a default value, physical units, a description, and optional ontology identifiers (GO [22], ChEBI [23], SBO [24]). This metadata is used for composition validation and gives an automated agent a machine-readable account of what each connection means and whether two ports intended to share a variable are congruent. The topology wiring is a map from each process’s port names to shared state paths. Topology is held outside the processes themselves so the same process can be wired into multiple different composites, or have one input redirected to another model’s output, without editing model code. The Store holds the shared state as a flat vector for computational efficiency and exposes a dictionary interface during construction. A Composite bundles a set of processes with a topology and compiles the whole into a single ODE system right-hand side to be integrated via diffrax [25]. Composites themselves are composable, so hierarchies of published models can be built through the same mechanism. At composition time, the validation layer checks and, where possible, reconciles units through the pint quantity system [26]. Ontology annotations are compared by identifier, flagging a conflict for any incompatible ontology terms. Additionally, a graph analysis of the wired topology reports highly connected nodes and overall coupling density, prompting revision of potentially unintended topologies.

### 2.2 Scheduler

Biological subsystems rarely share a clock: a signaling cascade may reach equilibrium in seconds, a transcriptional program in hours, and mitochondrial and DNA-damage dynamics over days or longer. When such processes are integrated as one coupled system, the fastest component dictates the step size for all of them, and a single stiff reaction can force the whole composite down to minuscule steps. This is a major obstacle to multi-scale mechanistic simulation, and it is the problem the Scheduler exists to solve.

Given a composite and a time span, the scheduler integrates the coupled system and returns the state over time. This single-runner design follows the *director* abstraction of Ptolemy II [27], in which one component governs how a heterogeneous collection of subsystems is scheduled and advanced in time. Its design goal is to keep the fastest or stiffest process from penalizing the rest through timescale grouping. When a composite occupies a single timescale, the Scheduler hands the whole system to one adaptive diffrax integration across the full interval. When processes span several timescales, they are automatically partitioned into groups whose characteristic rates fall within a bounded ratio, and each group is integrated independently by operator splitting [28]. Over a short window, one group is integrated while the others are held as input, and the groups exchange their updated states before the next macro window. Discrete and event processes contribute at the window boundaries as well. Splitting is first order Lie by default, where each group is advanced once per window, but it can also be symmetric Strang, where a group is advanced over two mirrored half-windows around the others. Within a window a group sees the others either as a frozen snapshot taken at the window start or as a piecewise-linear interpolant of their trajectory across the window. The interpolant exists only for groups already advanced in that window, so it is used automatically wherever a group reads a variable of one solved before it. Since this leaves a gap for feedback loops, where reading in the other direction sees the frozen snapshot, we also implement windowed waveform relaxation [29], closing such a loop by re-solving the window for a fixed number of sweeps, each group reading the others’ trajectory from the previous one.

The Scheduler selects a solver for each group separately, by estimating the fastest relaxation rate from the group’s Jacobian. If an explicit method would require an impractical number of substeps, that group is routed to an implicit solver (Kvaerno5), while non-stiff groups keep a cheaper explicit method (Tsit5). For large groups the fastest rates are estimated from a few hundred products of the Jacobian with trial vectors instead of the full Jacobian, staying memory-efficient and available at genome-scale group sizes (∼10^4^ states). The whole multigroup integration is just-in-time (JIT) compiled and differentiable, so a gradient defined on any downstream readout propagates back through the integration to any upstream parameter.

### 2.3 Model import and discovery

Much of the mechanistic corpus relevant to aging is distributed in SBML format, so cross-publication composition hinges on turning an SBML file into a Process. Similarly to SBMLtoODEjax [11] and jaxkineticmodel [12], we import an SBML file and transcribe its species and reactions into a JAX right-hand side. We wrap its output as a Process, readable and writable by other processes. Composites can in turn be exported as SBML. The framework also supports import of XPP and COPASI files, allows custom models written in Python, and has templates for composing and training Neural ODE models.

To discover new models, hallsim searches BioModels, JWS Online, ModelDB, BioSimulations, Physiome and Europe PMC, recovering models that were either deposited in popular model repositories or exist as supplementary files for a scientific paper. If a paper links its code instead of depositing a model, the framework follows the code-availability statement to GitHub, GitLab or Bitbucket. Another search over GEO and Zenodo finds potential datasets to calibrate against.

Each candidate is first evaluated on its own before composition. A first pass triage rejects models that fail to import or carry unsupported constructs, flagging potential concerns such as tolerance-sensitivity, a vanishing trajectory, a non-finite gradient, or a missing time unit. If the original paper deposits its fitting data, the pre-intake screen also asks whether the deposit still reproduces the fit of its publication. Models that pass the mechanical screen can then be reviewed in depth by role-specific LLM prompts, to evaluate both the biological common sense and mathematical validity of the models and the resulting composites.

### 2.4 Perturbation handles

A perturbation handle is a severity attached to a set of mappings, each pointing at a mechanistic parameter in a process and specifying how that parameter moves as severity changes. The hallmarks of aging provide one application of these handles. While the hallmarks are not orthogonal phenomena, it is difficult to visualize how perturbing one hallmark affects the others. Hallsim represents each hallmark as a perturbation handle and allows moving its severity to observe the downstream effects, which could serve as an educational tool, supported by hallsim’s primitives for building a visualizer web app. For instance, a Genomic Instability handle drives the DNA-damage input, Deregulated Nutrient Sensing acts on mTOR phosphorylation rate. An experimental arm (untreated disease, drug rescue, healthy control) can be modelled as a choice of severities over the same underlying composite, where the experimental condition and a pharmacological intervention are expressed on the same axis. Severity is a differentiable input which also allows for analysis of model sensitivity to a hallmark.

## 3. Results

### 3.1 Cross-publication composability verification

We demonstrate compositional co-simulation and joint calibration on a composite that stitches three independently published SBML models (Table 1) and four custom coupling edges into a single differentiable system conceptually spanning four hallmarks of aging (Figure 1). The composite exposes two control inputs representing genomic instability and deregulated nutrient sensing. We compare how the model behaves in control and perturbed conditions.

**Table 1.** Independently developed models merged into a composite.

|  | Source (BioModels) | Native clock | Axis | Total parameters | Integrated species |
| --- | --- | --- | --- | --- | --- |
| dp14 | Dalle Pezze et al (2014)[30]<br>BIOMD0000000582 | days | Nutrient-sensing,<br>senescence,<br>mitochondrial dysfunction | 56 | 23 |
| p07 | Proctor et al (2007)[31]<br>BIOMD0000000105 | seconds | Proteostasis | 22 | 35 |
| gz06 | Geva-Zatorsky et al (2006)[32]<br>BIOMD0000000157 | hours | Genomic instability | 9 | 3 |

**Figure 1.**
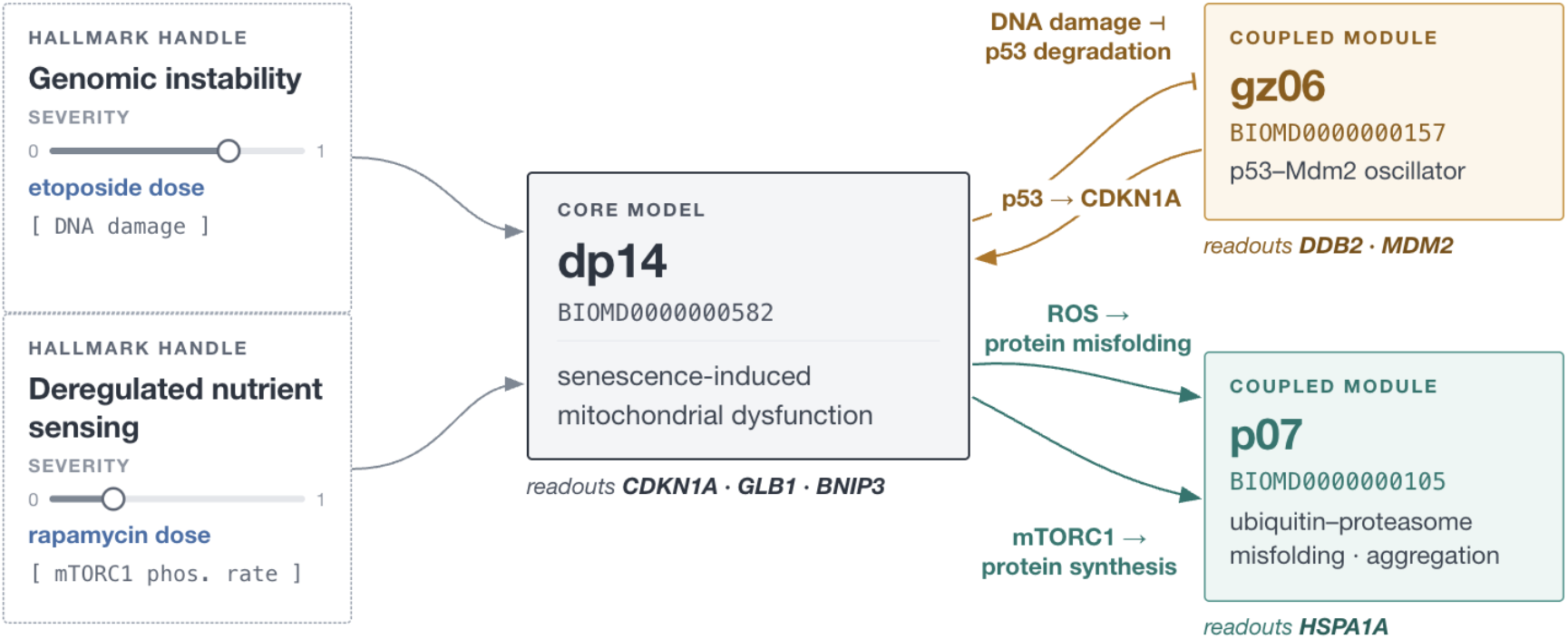
An example composite built by combining three independently published models through four coupling edges

The initial model choice was grounded in the work of Kounis et al [4] who surveyed models of cellular dynamics relevant for aging. The central model, referred here as dp14, is a cellular-senescence network model [30] that dynamically couples DNA-damage signaling, insulin– mTOR nutrient sensing, FoxO3a, oxidative-stress response, and mitochondrial turnover. The second model (p07) is a stochastic proteostasis model [31]. We chose to omit the third AMPK/mTOR model selected by Kounis, because it largely overlaps the species already inside the main senescence model. Instead, we extend the composite through a simple yet foundational p53–Mdm2 oscillator (gz06) [32] for the explicit DNA-damage-response hub, which the senescence model abstracts. The three models declare different native timescales of days (dp14), hours (gz06) and seconds (p07) which the framework rescales onto a canonical one-unit-per-day clock, and auto-clusters the processes into two timescale groups solved by operator splitting. We found that operator splitting on this composite runs 2.7× faster than a single implicit solve of the whole system at the same tolerances, and that the splitting benefit grows with the size of the model, reaching up to 30x speed-up on a synthetic 10,000 state model.

Dp14’s accumulated DNA damage was connected to suppress Geva-Zatorsky’s Mdm2- independent p53 degradation, abstracting away phosphorylation of ATM as the intermediate effector [33]. The gate is placed so that the etoposide treated arm crosses the oscillator’s bifurcation and the control arm does not. In the other direction, the p53 level drives p21 transcription in dp14 [34] through a saturating Hill term. Dp14’s ROS pool is read by p07’s misfolding law through a linear gain that converts between the two models’ concentration scales, and dp14’s phosphorylated mTORC1 sets p07’s protein-synthesis rate through a linear map, anchored at two points: the p07 published rate at the reference mTORC1 level, and the roughly halved rate under complete mTORC1 inhibition [35]. The same four couplings run in every experimental arm, and each of their parameters is a differentiable quantity that can be fitted alongside the parameters inside the models.

We selected six reporters spanning all three constituent models and their associated aging processes: CDKN1A (dp14/p21) and GLB1 (dp14/SA-β-gal) for cell-cycle arrest and senescence, BNIP3 (dp14/FoxO3a) for the FoxO3a-driven mitophagy branch, DDB2 (gz06/p53) and MDM2 (gz06/Mdm2 transcript) for the p53 DNA-damage response, and HSPA1A (p07/misfolded protein) for proteostasis. Together they represent the senescence, mitochondrial, genomic-instability, and proteostasis branches of the model, to assess concordance across the whole coupled network. The two p53 targets, DDB2 and MDM2, are read as the root-mean-square (RMS) of their p53-driven signal, smoothed over a few pulses and computed post hoc on the solved trajectory. This attenuates the sampling phase dependence, and accounts for the fact that in the gz06 model the mean p53 is not sensitive to damage, but the pulse amplitude is. The reporter levels are expressed as log2 fold change from baseline.

The simulated behavior is compared to the measured one using GSE248823, a public dataset of human fibroblasts undergoing etoposide-induced senescence [36]. The dataset provides bulk microarray expression at baseline (day 0), day 7, and day 14 in two biological replicates per arm. The different arms include etoposide DNA-damage induced senescence (DDIS) where etoposide pulse was applied over two days, etoposide with rapamycin rescue (RAPA) where rapamycin intervention was introduced at day two after the etoposide exposure ended, and an oncogene-induced stress experimental arm which we didn’t use. This dataset was chosen for demonstration based on its accessibility (public GEO portal), topical alignment (gerotherapeutic modulation of senescence), and perturbation and rescue design.

We demonstrate the composite’s end-to-end differentiability across the joined models, by unfreezing six mechanism parameters that lie along the routes from intervention handles to gene readouts and calibrating them to the data. The dataset’s three time-point trajectory is a rather sparse time course that constrains rather than fully resolves the underlying dynamics, and the framework’s identifiability analysis suggested freezing two of those parameters as unidentifiable from data. The parameters we tuned thus include: dp14’s CDKN1A transcription, DNA repair and SA-β-gal decay, as well as gz06’s MDM2 degradation. The loss used is the mean of squared errors between the model and data log2FCs over all reporter × timepoint pairs, and an added regularization term to penalize large deviations from the published parameter values. The parameters are optimized by gradient descent using Adam optimizer [37]. Calibration used only the DDIS-vs-control arm, while the rapamycin (RAPA) arm was held out and scored afterwards as a test of generalization.

The calibration loss decreased monotonically over 150 epochs and the mean reporter error against the experimental data fell on both the training and held out arms (Table 2). Figure 2. shows the species the six reporters read in the control, etoposide and etoposide-plus-rapamycin arms after calibration. From day 2 at which the rapamycin intervention was introduced, the rapamycin arm sits below the etoposide arm in all reporters. The control arm behavior is the opposite than what is expected in dp14 reporters GLB1 and BNIP3, due to the original article not modelling for a control state at all. GLB1 saw the largest movement towards the data measurements (Figure 3), while BNIP3 moved slightly in the opposite direction, since no free parameter was left on its path that could act as a lever.

**Table 2.** Calibrated vs out-of-the-box model-data concordance in mean absolute error.

|  | Day | MAE<br>*published* | MAE<br>*calibrated* |
| --- | --- | --- | --- |
| *DDIS (fit)* | 7 | 0.35 | 0.29 |
| *DDIS (fit)* | 14 | 0.42 | 0.27 |
| *RAPA (held-out)* | 7 | 0.29 | 0.23 |
| *RAPA (held-out)* | 14 | 0.40 | 0.27 |

**Figure 2.**
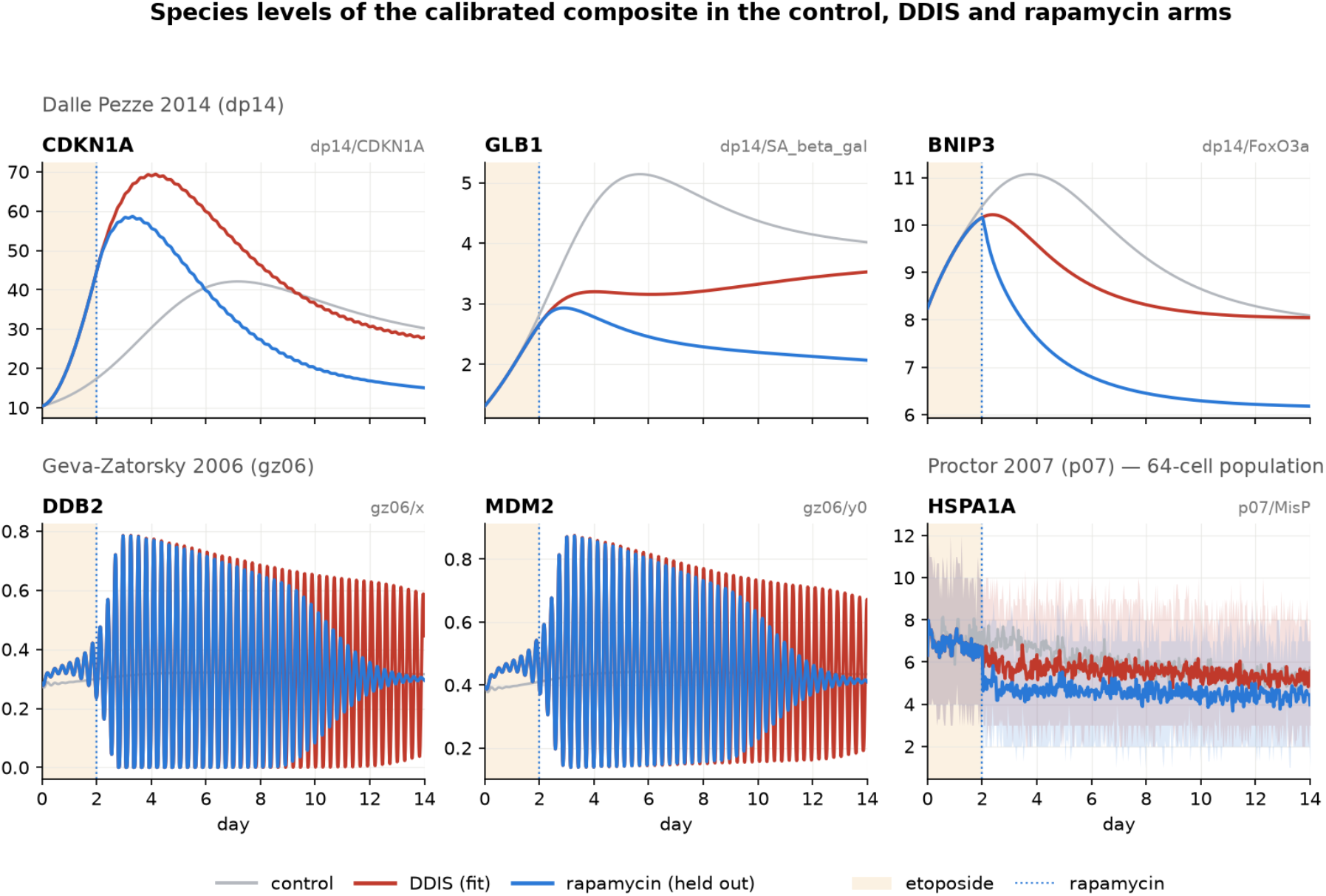
Species trajectories of the calibrated composite in control, DNA Damage Induced Senescence (DDIS) and rapamycin-treated arms.

**Figure 3.**
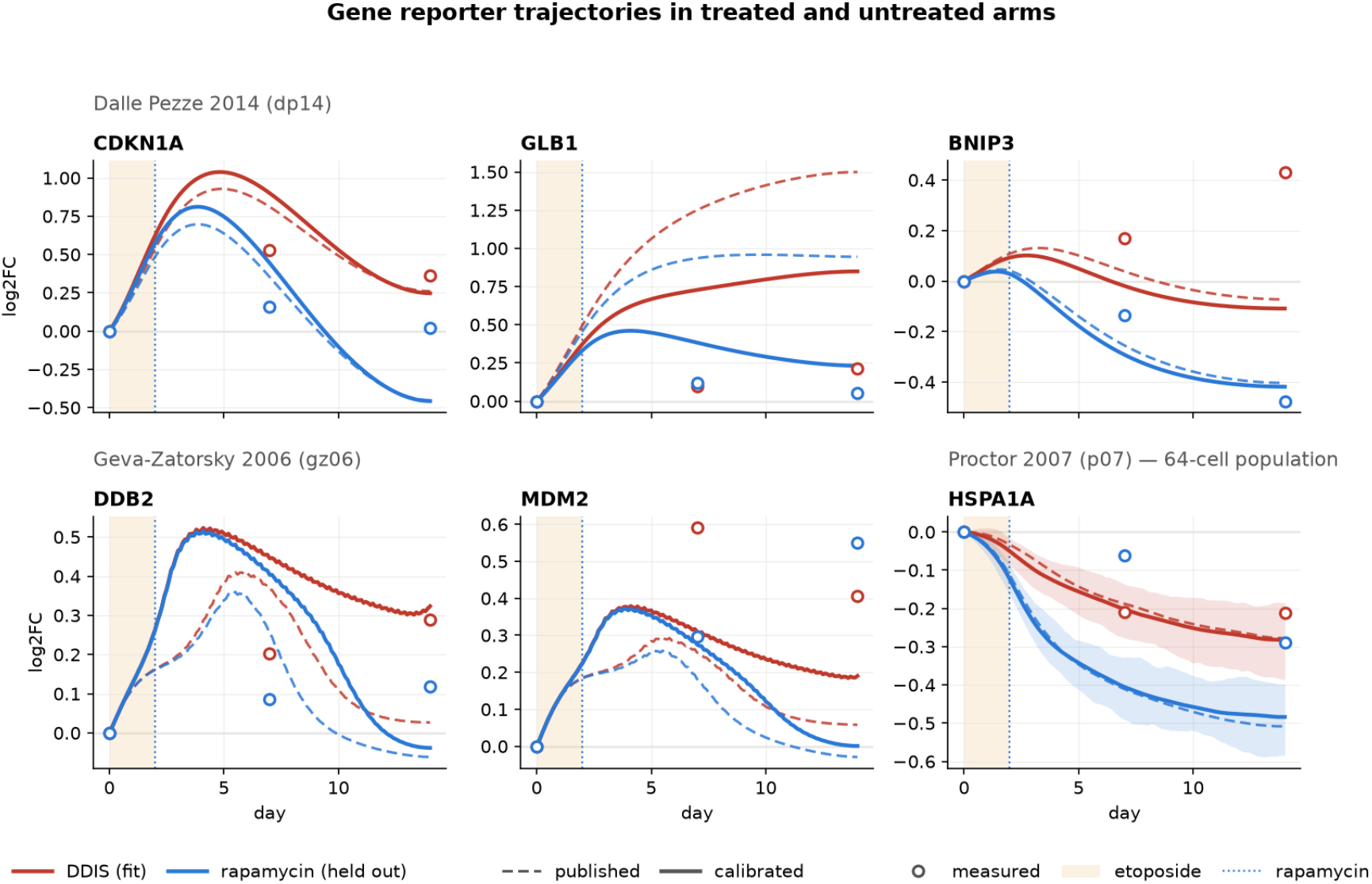
Temporal trajectories for six gene reporters in DNA Damage Induced Senescence (DDIS) and rapamycin-treated arms expressed in log2 fold change compared to baseline for published parameters and calibrated composite models.

Aside from the original dp14 model having no control arm, we also observed that its starting state is not in equilibrium. The cell spontaneously senesces, even without the DNA damage stimulus. The control and DNA damage arms converge to the same attractor, and the damage pulse does not affect the vector field. We also noticed that p07 models a ubiquitin pool with no synthesis and only degradation as exit, so a rapamycin treatment that lowers misfolding leaves ubiquitin charged on the conjugating enzymes and drains the free pool. More work on aging models is thus required to properly assess intervention potential. A more complete composite also needs aggregate clearance wired to autophagy through mTORC1, ULK1 and p62, more work on how single cell p53 protein pulsing translates to downstream bulk RNA readouts, and a late-rising inflammatory module with NF-κB, IL-6 and JAK/STAT [38], [39], [40].

### 3.2 Population batching

The spread across a cell population and pooled effects are usually of interest more than a single cell trajectory. Population analyses are simple in hallsim, which expects a batch axis with varying initial conditions, running through every group’s solve as one batched computation, with no per-cell loop. Cells that interact, by exchanged signals or in a tissue, lie beyond the scope of the independent-cell models presented here.

The p53–Mdm2 oscillator in the gz06 part of the composite comes from a study [32] whose central finding is the cell-to-cell variability of the response. Simulating the oscillator as a population with lognormal spread in its rates shows the heterogeneity that a bulk assay sees (Figure 4). Spread in the production rate changes amplitudes and leaves the population mean oscillating coherently, while the spread in the Mdm2 degradation rate changes periods and dephases the cells, resulting in a damped oscillation. Adding random initial phase attenuates the bulk amplitude further.

**Figure 4.**
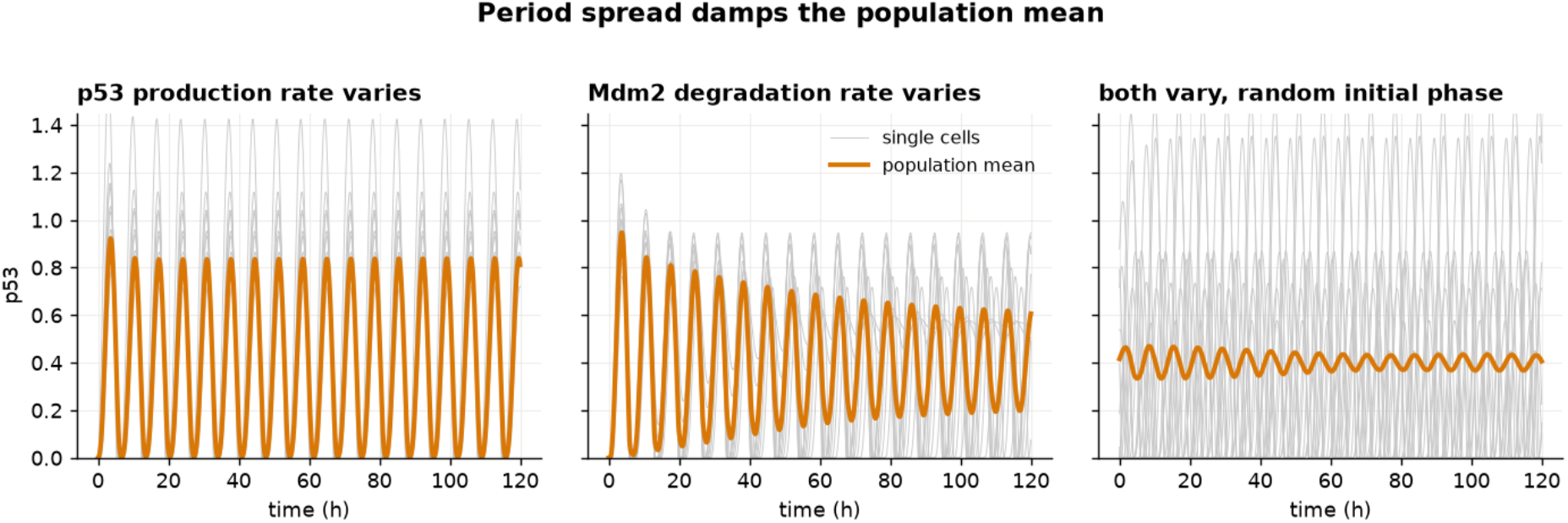
Simulating batch populations shows how sustained oscillations at the cell level produce a damped oscillator in bulk.

### 3.3 Hybrid composition

To show that mechanistic and learned components compose interchangeably, we train a surrogate and attach it in place of one mechanistic sub-model. The gz06 block’s right-hand side is replaced by a Neural ODE process, a multilayer perceptron with three hidden layers of 192 units mapping the block’s state and its two conditioning inputs to a time derivative. The conditioning inputs are the p53 degradation rate α_x_, and the Mdm2 degradation rate α_y_. Training trajectories are simulated from the mechanistic model over a grid of both rates and a range of initial states, so the network learns a family of vector fields instead of one. It is fit by derivative matching, then refined by multiple shooting [41] with a collocation term [42]. The model crosses two Hopf bifurcations as α_y_ varies, from a fixed point through a limit cycle to damped oscillation, and one as α_x_ varies. The surrogate reproduces all three regimes (Figure 5, top) and recovers the pulse amplitude across the α_y_ sweep including its two bifurcation points (Figure 5, bottom left). The one visible discrepancy is a small transient in the fixed-point regime near the lower Hopf. Composed back into the composite beside the untouched dp14 and p07 blocks, the hybrid reproduces the mechanistic DDB2 readout of p53 across genomic-instability severity (Figure 5, bottom right), and the gradient of that readout with respect to severity passes through the neural block, agreeing with a finite difference. The learned process is differentiable and composable like a mechanistic one.

**Figure 5.**
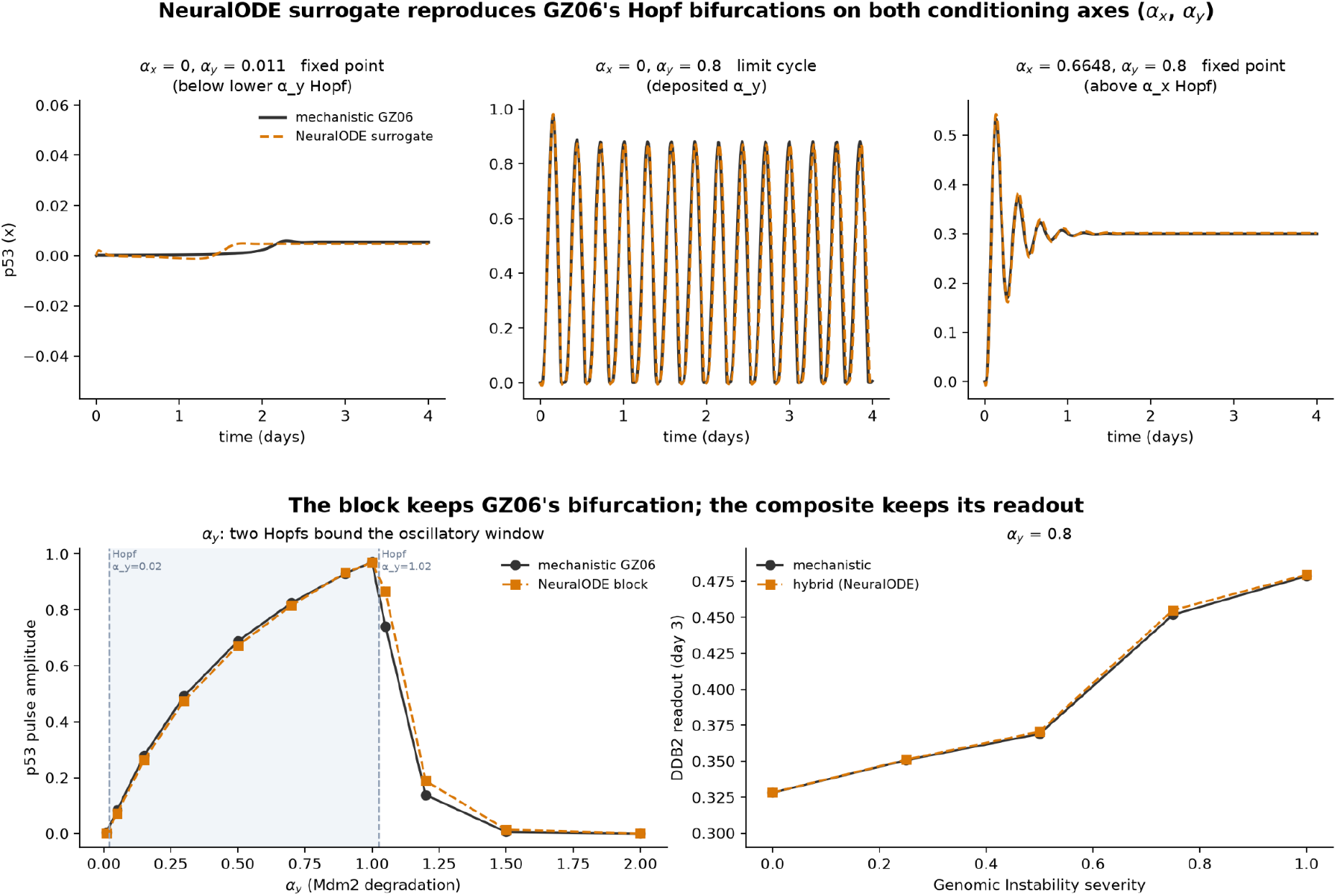
A Neural ODE surrogate of the GZ06 p53–Mdm2 oscillator, conditioned on the p53 degradation rate α_x_ and the Mdm2 degradation rate α_y_. Top: p53 over time, mechanistic against surrogate, in the three regimes; the α_y_ values here are outside the training grid.Bottom left: pulse amplitude across α_y_ with the two bifurcation points. Bottom right: the composite’s DDB2 readout across genomic-instability severity with the mechanistic block and with the surrogate in its place.

## 4. Discussion

We developed a JAX-native framework for composing independently published kinetic models into a single object that is end-to-end differentiable and executable as a batched population on a CPU or GPU. We enrich the framework with a number of automatic checks, guards and suggestions, in order to steer the user or AI agent towards building a composite without falling into common traps, envisioning it as the tool which will eventually help us construct a comprehensive model of cellular aging.

As a demonstration of capability, we composed three independently published models and four custom branches into a composite. From the technical point of view, composing a fourth model would be a matter of an import and a topology row, having found the model by using the framework’s methods for searching online model repositories and literature. We searched BioModels to gauge supply, and ran the pre-intake analysis over every deposited SBML file there. We found that out of the 2,531 total (1,075 curated and 1,456 uncurated) models, 1,567 run as published, and about 1,300 of them could be used for future composition, with minor tweaks such as establishing the native timescale, or annotating species. Other repositories can also be screened for potential sources of composable kinetic models. The framework’s validation layer examines the composite’s wiring and flags ontology mismatches and potentially incompatible connections for revision. The stiffness analyzer selects a solver for each group, and identifiability analysis highlights the parameters that the available data can constrain.

We show that calibration runs through the whole system and brings its predictions closer to the measured reporters on both the fitted and the held-out condition. An mTORC1 perturbation, represented as the hallmark Deregulated Nutrient Sensing, shows effects that are consistent with rapamycin ameliorating some of the consequences of an etoposide pulse (which is conceptually represented as an increase in Genomic Instability). We train a neural surrogate of a submodel, and swap it in place of its mechanistic parent, demonstrating that the neural component composes and runs just as a mechanistic one, while reproducing the output across a sweep of parameters and two bifurcations. While deep neural models tend to interpolate within the support of their data, mechanistic models can better extrapolate from causal structure, and the field has spent decades encoding that structure. We believe that the choice between mechanistic modelling and machine learning is a false dichotomy: a neural module can sit inside a composite, as we show here, and a mechanistic composite’s simulated state, including its state under an intervention, can be a feature for a deep learning model predicting organism level phenotypes.

The framework and the composition demonstration in this paper have several limitations. Hallsim currently focuses on ODE dynamics and spatial cell-to-cell communication is future work. While forward evaluation can run in single precision, calibration requires double precision 64bit floating point to support tight error tolerances of models with fast dynamics. This may underutilize certain types of GPUs for composite calibration. The data used for calibration of the demo composite is bulk microarray, a destructive assay that doesn’t allow pairing across timepoints but only estimates the population mean. Additionally, with two replicates per condition and three available timepoints, the data are insufficient to properly constrain a trajectory, but instead just mildly steer it. This modelling exercise also revealed unreported limitations of the original unit models themselves, which call for careful future revisions of this core aging biology composite.

## 5. Data and code availability

The full code is open sourced at https://github.com/BabaJaguska/hallsim.git. The dataset used for demonstration in this paper was downloaded from GEO under the accession number GSE248823 (https://www.ncbi.nlm.nih.gov/geo/query/acc.cgi?acc=GSE248823). The composed and calibrated model is deposited in BioModels under identifier MODEL2609140001.

## 6. Acknowledgements

We thank Marko Milovanovic for early review and brainstorming.

## 8. Author contributions

M.B. and D.F conceptualized the framework and wrote the paper. M.B. designed and implemented the computational solution.

